# Effect of number and placement of EEG electrodes on measurement of neural tracking of speech

**DOI:** 10.1101/800979

**Authors:** Jair Montoya-Martínez, Jonas Vanthornhout, Alexander Bertrand, Tom Francart

## Abstract

Measurement of neural tracking of natural running speech from the electroencephalogram (EEG) is an increasingly popular method in auditory neuroscience and has applications in audiology. The method involves decoding the envelope of the speech signal from the EEG signal, and calculating the correlation with the envelope of the audio stream that was presented to the subject. Typically EEG systems with 64 or more electrodes are used. However, in practical applications, set-ups with fewer electrodes are required. Here, we determine the optimal number of electrodes, and the best position to place a limited number of electrodes on the scalp. We propose a channel selection strategy based on an utility metric, which allows a quick quantitative assessment of the influence of a channel (or a group of channels) on the reconstruction error. We consider two use cases: a subject-specific case, where the optimal number and position of the electrodes is determined for each subject individually, and a subject-independent case, where the electrodes are placed at the same positions (in the 10-20 system) for all the subjects. We evaluated our approach using 64-channel EEG data from 90 subjects. In the subject-specific case we found that the correlation between actual and reconstructed envelope first increased with decreasing number of electrodes, with an optimum at around 20 electrodes, yielding 29% higher correlations using the optimal number of electrodes compared to all electrodes. This means that our strategy of removing electrodes can be used to improve the correlation metric in high-density EEG recordings. In the subject-independent case, we obtained a stable decoding performance when decreasing from 64 to 22 channels. When the number of channels was further decreased, the correlation decreased. For a maximal decrease in correlation of 10%, 32 well-placed electrodes were sufficient in 91% of the subjects.

## Introduction

To understand how the human brain processes an auditory stimulus, it is essential to use ecologically valid stimuli. An increasingly popular method is to measure neural tracking of natural running speech from the electroencephalogram (EEG). This method also has applications in domains such as audiology, as part of an objective measure of speech intelligibility (Vanthornhout et al., 2018; Lesenfants et al., 2019), and coma science (Braiman et al., 2018).

Among the different models used to study the relationship between the stimulus and the brain response, two of the most often used ones are the forward and backward models (e.g., Crosse et al., 2016; Lalor and Foxe, 2010; Ding and Simon, 2012; Verschueren et al., 2019; Vanthornhout et al., 2018). In the forward model (also know as encoding model), we determine a linear mapping from the stimulus to the brain response. In the backward model (also known as stimulus reconstruction), we determine the linear mapping from the brain response to the stimulus. Backward models are referred to as decoding models, because they attempt to reverse the data generation process. Both the forward and backward models involve the solution of a linear least squares (LS) regression problem. The quality of the reconstruction is usually quantified in terms of correlation between the true and reconstructed signal. The benefit of the forward model is that the obtained models (also called temporal response functions) can be easily interpreted, and topographical information can be easily obtained. The benefit of the backward model is that through combination of information across EEG channels, better performance (higher correlations) can be obtained, but the model coefficients can not be easily interpreted. Another approach to study the relationship between the stimulus and the brain response is based on Canonical Correlation Analysis (CCA) (Hotelling, 1936). CCA estimates the optimal linear operator to be applied to both the stimulus and the response in order to reveal correlations between the two. This allows the stimulus representation to be stripped of dimensions irrelevant for measurable brain responses, and the EEG to be stripped of activity unrelated to auditory perception (de Cheveigné et al., 2018). In this experimental paradigm, the most used stimulus representation is its slowly varying temporal envelope (e.g., Ding and Simon, 2011; Aiken and Picton, 2008), which is known to be one of the most important cues for speech recognition (Shannon et al., 1995).

EEG systems used in research typically have 64 electrodes or more. However, in practical applications, such as objective measurement of speech intelligibility in the clinic, such large numbers of electrodes are not always possible due to the cost of high density systems and the time required to place the electrodes on the scalp. We therefore considered the following questions: for a smaller number of electrodes, (1) what is the optimal location of electrodes on the scalp and (2) what is the impact on the correlation when we decrease the number of channels. We consider two use cases: in one case the electrodes are placed at the same positions (in the 10-20 system) for all subjects, which would for instance be relevant in the design of an application-specific headset or electrode cap. We will refer to this use case as the *subject-independent scenario*. In a second use case the optimal number and position of electrodes is determined for each subject individually. In this case the main application of the method would be to improve correlations with the use of a high-density system. Another future application would be to determine the optimal electrode locations in order to design a custom system for each subject, but this will require further validation. We will refer to this use case as the *subject-specific scenario*. We use the backward model, due to its advantages in decoding accuracy compared to the forward model.

We started from 64-channel recordings, and considered the question which subset of *K* channels allow to get the best decoding performance. This is a combinatorial problem, closely related to the column subset selection problem (Boutsidis et al., 2009), whose NP-hardness is an interesting open problem. In order to overcome this challenge, Mirkovic et al. (2015); Fuglsang et al. (2017) used a channel selection strategy based on an iterative backward elimination approach, where at each iteration, the electrode with the lowest corresponding coefficient magnitudes in the decoder is removed from the next iteration (we will refer to this channel selection method as the decoder magnitude-based (DMB) method). This strategy assumes that important channels will have a large coefficient in the least squares solution. However, as pointed out by (Bertrand, 2018), this is an unsuitable assumption: for example, if the coefficients of one of the channels would all be scaled with a factor *α*, then the corresponding decoder coefficient in the LS solution would be scaled with *α*^−1^, whereas the information content of that channel obviously remains unchanged.

In this work, we propose a channel selection strategy based on the utility metric (Bertrand, 2018), which allows a quick quantitative assessment of the influence of a channel (or a group of channels) on the reconstruction error. In the subject-specific scenario, we use the utility metric to remove channels one by one, each time removing the channel with least influence on the reconstruction error. In the subject-independent scenario, we use the utility metric to remove one group of channels at a time (the group of channels with least influence on the reconstruction error). For channels located off the midline each group is composed of one channel located over the left hemisphere and its closest symmetric counterpart located over the right hemisphere. For channels located over the central line dividing both hemispheres, each group is composed of one channel located over the frontal lobe and its closest symmetric counterpart located either over the parietal or the occipital lobe (see Fig. 1). The rationale behind this channel selection strategy is to maintain symmetry. The symmetry criterion avoids bias to one hemisphere, which could be problematic as hemispheric differences are often found between subjects (e.g., Goossens et al., 2019; Van Eeckhoutte et al., 2018; Poelmans et al., 2012; Vanvooren et al., 2015). A similar channel selection strategy, also based on the utility metric, was proposed by Narayanan and Bertrand (2019) on an auditory attention decoding task, where the main goal was to optimize the topology of a wireless EEG sensor network (WESN), without imposing a symmetry constraint on the selected channels. We evaluated our approach using EEG data from 90 subjects. We aimed to minimize reconstruction error, and to minimize the intra-subject variability in reconstruction error.

**Fig 1.**
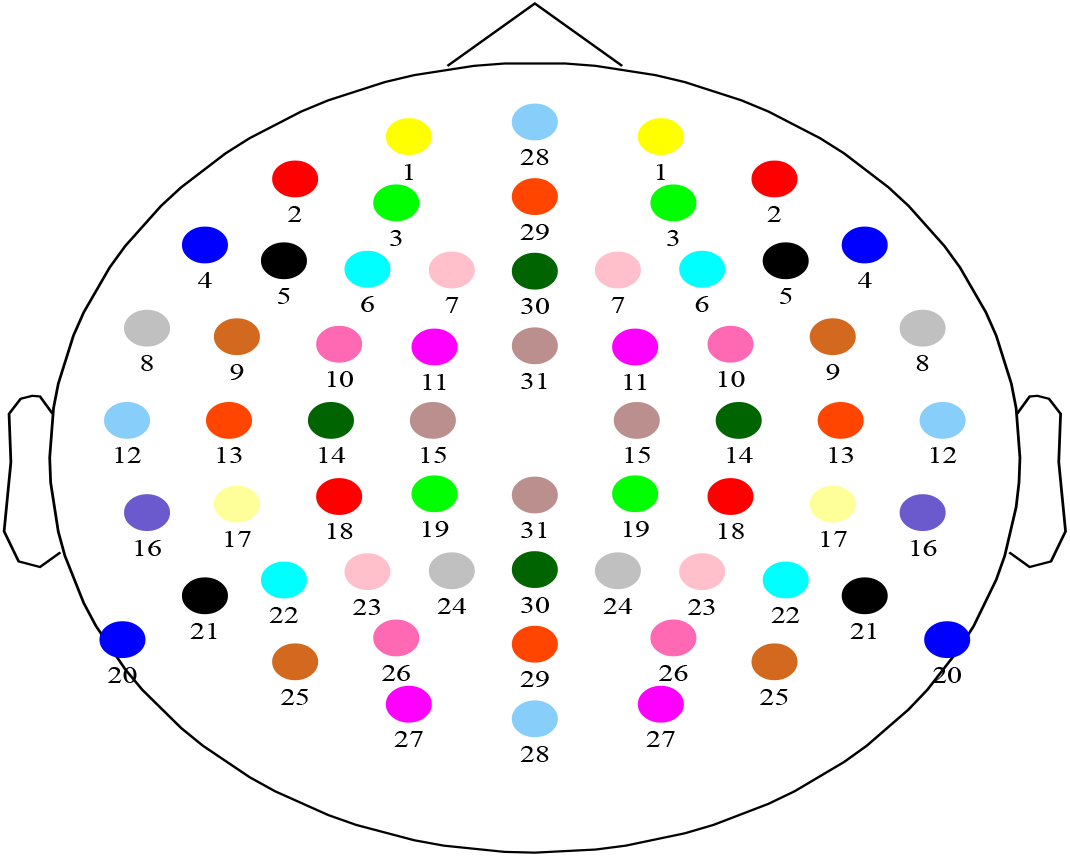
Channel grouping strategy. For channels located either over the left or right hemisphere (groups 1, 2, …, 27), each group is composed by one channel located over the left hemisphere and its closest symmetric counterpart located over the right hemisphere. For channels located over the central line dividing both hemispheres (groups 28, 29, 30, 31), each group is composed by one channels located over the frontal lobe and its closest symmetric counterpart located either over the parietal or the occipital lobe.

While Narayanan and Bertrand (2019); Mirkovic et al. (2015) used auditory attention decoding accuracy as their main outcome measure, we investigated the underlying correlation between actual and reconstructed envelope instead. This correlation can be considered a measure of signal to noise ratio. Note that stable attention decoding accuracy does not imply stable correlation. Another difference with the literature is that we considered both the subject-specific and subject-independent cases. In addition, we propose and use symmetry constraints for avoiding lateralization bias.

## Methods

### Data collection

#### Participants

Ninety Flemish-speaking volunteers participated in this study. Their age ranged from 18 to 30 years old, with a mean of 22. 15% of the participants was male and 7% was left-handed. They were recruited from our university student population to ensure normal language processing and cognitive function. No tests of language processing or cognitive function were performed. Each participant reported normal hearing, which was verified by pure tone audiometry (thresholds lower than 25 dB HL for 125 Hz until 8000 Hz using MADSEN Orbiter 922–2 audiometer). Before each experiment, the participants signed an informed consent form. The study was approved by the Medical Ethics Committee UZ KU Leuven/Research (KU Leuven).

#### Experiment

Each participant listened attentively to the children’s story “Milan”, written and narrated in Flemish by Stijn Vranken. The stimulus was 15 minutes long and was presented binaurally at 60 dBA without any noise. The participants were motivated to pay attention by asking content-related questions after presentation of the story (which were answered mostly correctly). It was presented through Etymotic ER-3A insert phones (Etymotic Research, Inc., IL, USA) which were electromagnetically shielded using CFL2 boxes from Perancea Ltd. (London, UK). The acoustic system was calibrated using a 2-cm^3^ coupler of the artificial ear (Brüel & Kjær, type 4192). The experimenter sat outside the room and presented the stimulus using the APEX 3 (version 3.1) software platform developed at ExpORL (Dept. Neurosciences, KU Leuven, Belgium) (Francart et al., 2008) and a RME Multiface II sound card (RME, Haimhausen, Germany) connected to a laptop. The experiments took place in a soundproof, electromagnetically shielded room.

#### EEG acquisition

In order to measure the EEG responses, we used a BioSemi (Amsterdam, Netherlands) ActiveTwo EEG setup with 64 channels. The signals were recorded at a sampling rate of 8192 Hz, using the ActiView software provided by BioSemi. The electrodes were placed over the scalp according to the international 10-20 standard.

### Signal processing

#### EEG pre-processing

Data pre-processing was performed offline, using MATLAB (Mathworks Inc., Natick, MA). In order to decrease computation time, the EEG data was downsampled from 8192 Hz to 1024 Hz (we used the *resample* function, which applies an antialiasing FIR lowpass filter to the data). Next, the data was bandpass filtered between 0.5-4 Hz (delta band), using a Chebyshev filter with 80 dB attenuation at 10 % outside the passband. Finally, the data was downsampled to 64 Hz and re-referenced to Cz in the channel subset selection stage, and to a common-average reference (across the selected channels) in the decoding performance evaluation stage. The delta band was chosen because it yields the highest correlations and most of the information in the stimulus envelope is contained within this frequency band (Vanthornhout et al., 2018; Ding and Simon, 2014). To assess the effect of frequency band on the results, we also analyzed the optimal number and placement of electrodes for measurement of neural tracking of speech in the theta band (4-8 Hz), the results are shown in the appendix.

The pre-processing pipeline does not include an artifact rejection step, as this would require the use of electrodes that may later on be eliminated and therefore can potentially leak information from the unselected channels to the selected ones. However, to investigate the effect of artefact rejection we repeated the full analysis using artifact rejection, involving the Sparse Time Artifact Removal method (STAR) (de Cheveigné, 2016) and the multi-channel Wiener filter algorithm (Somers et al., 2018). A Wilcoxon signed rank test of the effect of artefact rejection on correlation showed no significant effect.

#### Speech envelope

The speech envelope was computed according to Biesmans et al. (2017), who showed that good reconstruction accuracy can be achieved with a gammatone filterbank followed by a power law. We used a gammatone filterbank (Søndergaard et al., 2012; Søndergaard and Majdak, 2013), with 28 channels spaced by 1 equivalent rectangular bandwidth, with centre frequencies from 50 Hz to 5000 Hz. From each subband, we take the absolute value of each sample and raise it to the power of 0.6. The resulting 28 signals were then downsampled to 1024 Hz, averaged, bandpass filtered with a (0.5-4 Hz) Chebyshev filter to obtain the final envelope, and finally downsampled again to 64Hz. The power law was chosen as the human auditory system is not a linear system and compression is present in the system. The gammatone filterbank was chosen as it mimics the auditory filters present in the basilar membrane in the cochlea.

#### Backward model

The backward model to decode a speech envelope from the EEG can be stated as a regularized linear least squares (LS) problem (O’sullivan et al., 2014):

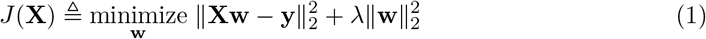

where **X** ∈ ℝ^*T*×(*N*×*τ*)^ is the EEG data matrix concatenated with *τ* time-shifted (zero-padded) version of itself, **y** ∈ ℝ^*T*×1^ is the speech envelope, **w** ∈ ℝ^(*N*×*τ*)×1^ is the decoder, *T* is the total number of time samples, *N* is the number of channels, *τ* is the number of time samples covering the time integration window of interest, and *λ* is a regularization parameter. The solution to the backward problem 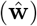 is usually referred to as a decoder. In order to choose the regularization parameter *λ*, we compute and sort the eigenvalues of the covariance matrix associated to **X**. Then, we pick as *λ* the eigenvalue where the accumulated percentage of explained variance is greater than 99%.

#### Channel selection

To select channels we used the utility metric (Bertrand, 2018), which quantifies the effective loss, i.e., the increase in the LS cost, if a group of columns (corresponding to one channel or a set of channels and all their *τ* − 1 corresponding time-shifted version) would be removed and if the model (1) would be reoptimized afterwards:

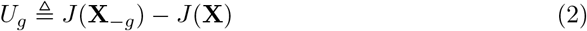

where **X**_*−g*_ denotes the EEG data matrix **X** after removing the columns associated with the *g*-th group of channels and their corresponding time-shifted versions. We will later on define how channels are grouped in our experiments (see Subsection).

Note that a naive implementation of computing *U*_*g*_ would require solving one LS squares problem like (1), for each possible removal of a candidate group, which would lead to a large computational cost for problems with large dimensions and/or involving a large number of groups.

Fortunately, this can be circumvented, as shown by Bertrand (2018), with a final computational complexity that scales linearly in the number of groups, given the solution of (1) when none of the channels are removed. The basic workflow for finding the best *k* groups of EEG channels can be summarized as follows (Narayanan and Bertrand, 2019): we compute the utility metric for each of the groups and remove the group with the lowest utility. Next, we recalculate the new values of the utility metric taking only into account the remaining groups, and once again we remove the one with the lowest value of utility. We continue iterating following these steps until we arrive to *k* groups.

We used the utility metric in two conditions: (1) in the subject-specific case where optimal electrodes are selected for each subject, and (2) in the generic case where the same set of electrodes is used for all subjects.

In the subject-specific case, we computed (for each subject *i*) the regularized covariance matrix 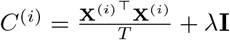 (**I** denotes the identity matrix) and the cross-correlation vector 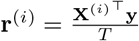 in order to compute the optimal all-channel decoder 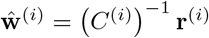. The utility metric for each (group of) channel(s) can be directly computed from 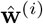 and *C*^(*i*)^ (we refer to Bertrand (2018) and Narayanan and Bertrand (2019) for more details). We used the utility metric toolbox from Narayanan and Bertrand (2019) available at https://github.com/mabhijithn/channelselect. We then ranked the groups according to their corresponding utilities, and removed the channel(s) corresponding to the group *g* with the lowest utility. We then repeated the same process with the matrix 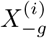 in which the columns corresponding to the channels in group *g* were removed. We kept repeating this process until only *k* groups remained.

Next, during the decoding evaluation stage, we computed a decoder by solving the backward problem using the best *k* selected groups of channels for each subject. In this stage, we re-referenced the channels with respect to the common average across the selected channels and discarded the reference electrode Cz. We solved each backward problem using a 7-fold cross-validation approach, where 6 folds were used for training and 1 for testing. This corresponds to approximately 12 and 2 minutes of data, respectively. This cross-validation served to reduce the influence of intra-suject variability over time on the results. Using the decoder 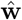, we computed the reconstructed envelope as 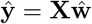 after which we computed the Spearman correlation between the reconstructed speech envelope 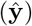 and the true one (**y**). By following this procedure, for each subject, we ended up with 7 values of correlation (corresponding to the evaluation of the correlation using each one of the test folds), which can be arranged as an array **S** ∈ ℝ^90×*k*×7^ (number of subjects × number of groups × number of test folds).

To compare with the literature, we also implemented the DMB approach, wherein we iteratively solved a backward problem for each subject, and at each iteration, the group of electrodes with the lowest corresponding coefficient magnitudes in the decoder was removed from the next iteration.

As a reference, we also implemented the forward model, where for each subject and electrode the correlation between actual EEG and EEG predicted from the speech envelope is obtained. The results are shown in the appendix.

In the subject-independent case, where the same set of electrodes is used for all subjects, we only used the utility metric. The evaluation consisted of the same two stages described above. The only difference was that, during the channel selection stage, we computed a grand average model by averaging the covariance matrices of all the subjects, which is equivalent to concatenating all the data from all the subjects in the data matrix *X* in (1). Finally, the decoding evaluation stage followed exactly the same steps described for the subject-specific case above, i.e., using a subject-*specific* decoder (yet, computed over electrodes that were selected in a subject-*independent* fashion).

#### Symmetric grouping of the EEG channels

In addition to selecting individual channels to remove (no grouping of channels), we also evaluated a strategy in which symmetric groups of channels were removed, to avoid hemisphere bias effects across subjects. Each group is composed of two EEG channels (see Fig. 1). For channels located on either side of the midline (Fig. 1, groups with labels from 1 to 27), each group is composed by one channel located over the left hemisphere and its closest symmetric counterpart located over the right hemisphere. For channels located over the midline dividing both hemispheres (Fig. 1, groups with labels from 28 to 31), each group is composed by one channel located over the frontal lobe and its closest symmetric counterpart located either over the parietal or the occipital lobe. Channel Cz does not belong to any group because it was used as a reference (in the channel subset selection stage). Channel Iz was not considered in order to preserve the symmetry with respect to the number of electrodes.

## Results

### Channel selection strategies: utility metric vs DMB

We compared the performance of the utility metric and DMB in the the subject-specific case, where the optimal electrode locations were determined for each subject individually. We compared the median of the correlation between **y** and 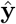 for each subject, as well as the number of channels required to obtain it (from now on referred to as the optimal number of channels). For both methods we observe a large increase in correlation when we use a reduced number of channels, with the optimum of the median around 20 and 30 channels, for the utility metric and DMB, respectively (see Fig. 2a). This means that the evaluated strategies of removing electrodes can be used to substantially improve the correlation metric in high-density EEG recordings.

**Fig 2.**
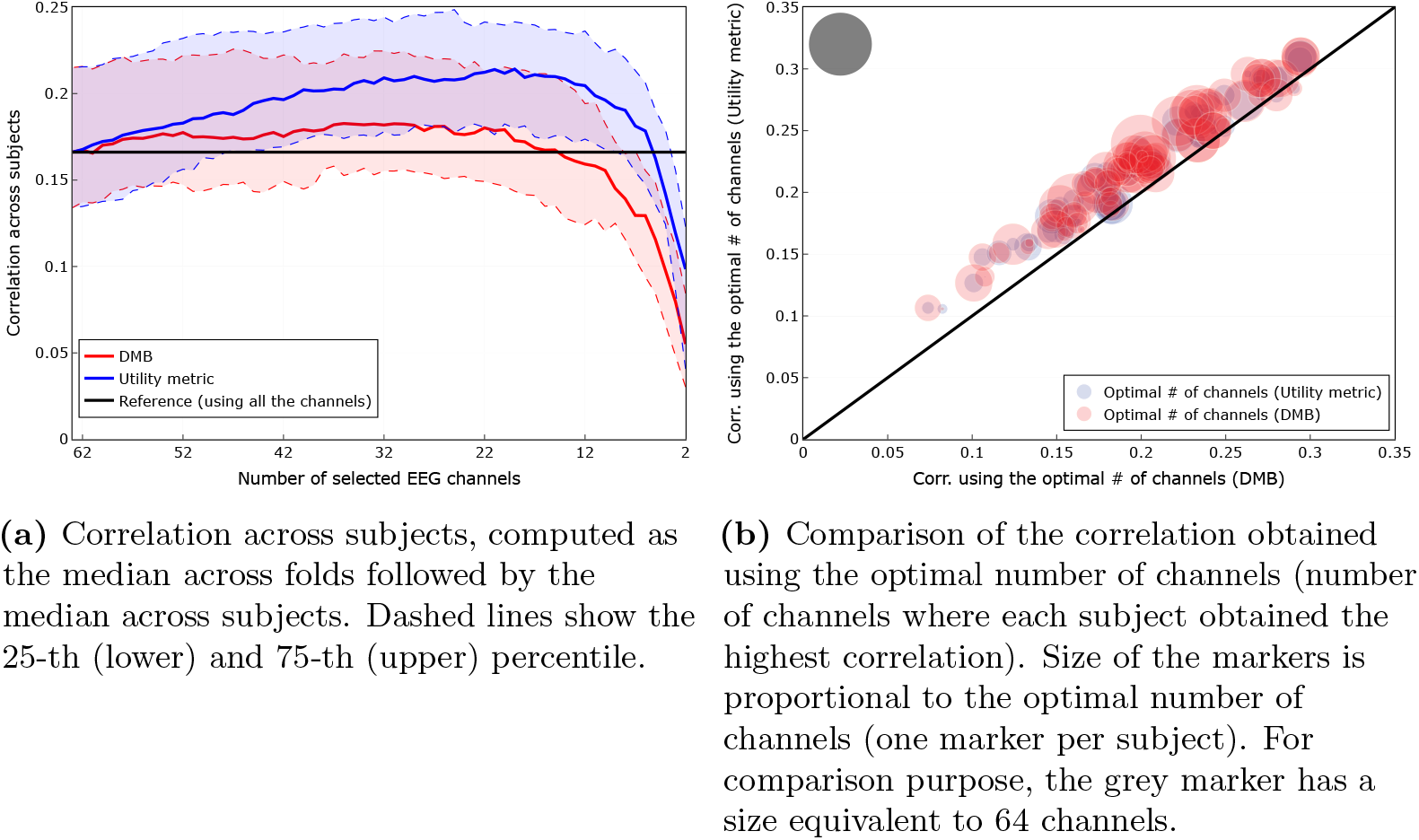
Comparison of channel selection strategies: utility metric vs DMB (*subject-specific scenario*). A Wilcoxon signed rank test showed that there was a significant difference (W=6, p ¡ 0.001) between the correlation obtained using the optimal number of channels according to the utility metric (median=0.22) compared to the one obtained using DMB (median=0.19). Another Wilcoxon signed rank test showed that there was also a significant difference (W=424.5, p ¡ 0.001) between the optimal number of channels selected by the utility metric (median=19) compared to the one selected by DMB (median=32).

We can see in Fig. 2a that the utility metric globally outperforms the DMB approach, obtaining consistently higher correlations (median) across subjects. In Fig. 2b, we can see that the utility metric also outperforms the DMB approach on an individual level, obtaining for every subject a higher value of maximal correlation, as well as requiring a smaller number of electrodes to obtain it. A Wilcoxon signed rank test showed that there was a significant difference (W=6, *p* < 0.001) between the correlation using the optimal number of channels according to the utility metric (median=0.22) compared to the one obtained using DMB (median=0.19). Another Wilcoxon signed rank test showed that there was also a significant difference (W=424.5, *p* < 0.001) between the optimal number of channels selected by the utility metric (median=19) compared to the optimal number selected by DMB (median=32). Because of the improved performance offered by the utility metric compared to DMB, we solely focus on the former in the remaining of the paper.

### Channel selection based on the utility metric vs using all the channels

In this section, we compare the channel selection strategy based on the utility metric with the case where all the available channels are used. We compared both strategies in the subject-specific and subject-independent scenario.

#### Subject-specific electrode locations

We consider here the condition where we remove channels one by one, obtaining the best channels for each subject independently. Fig. 3a shows the median correlation, computed as the median across folds followed by the median across subjects. Blue dashed lines show the 25-th (lower) and 75-th (upper) percentile. In this figure, we can see that at least 50% (median) of the subjects exhibit a higher correlation for 6 up to 63 channels, with respect to the correlation obtained with all the available channels.

**Fig 3.**
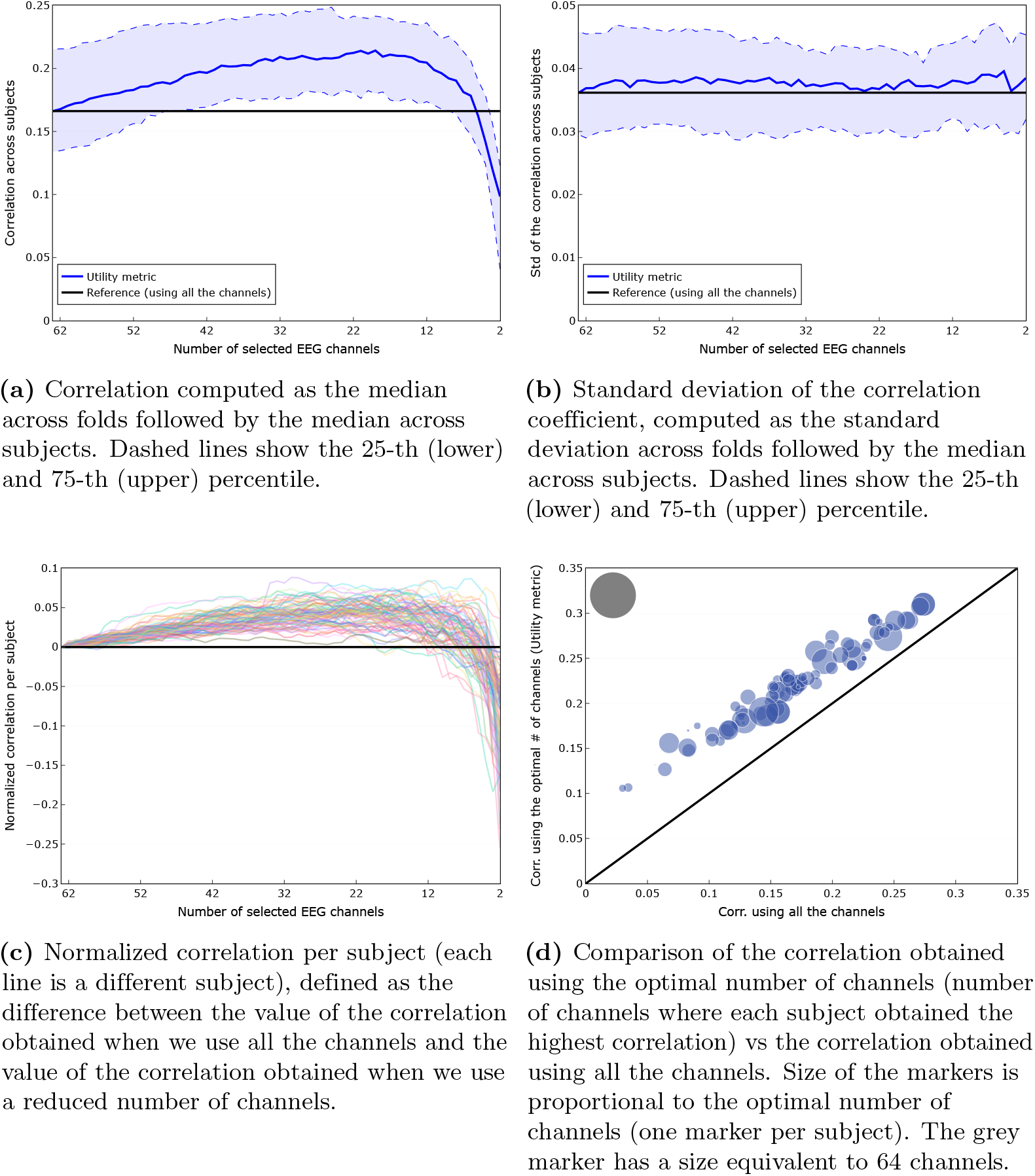
Comparison of the channel selection based on the utility metric vs using all the channels (*subject-specific scenario*). A Wilcoxon signed rank test showed that there was a significant difference (W=0, *p* < 0.001) between the correlation obtained using the optimal number of channels suggested by the utility metric (median=0.22) compared to the one obtained using all the channels (median=0.17).

Fig. 3b shows the standard deviation of the correlation, as a measure for within-subject variability, computed as the standard deviation across folds followed by the median across subjects. Blue dashed lines show the 25-th (lower) and 75-th (upper) percentile. In this figure we can see a largely stable standard deviation of the correlation around the reference value (standard deviation of the correlation when using all the 64 channels).

Figs. 3a and 3b suggest that we could obtain a higher correlation with a reduced number of channels. However, these are group results. Fig. 3c shows, independently for each subject, the difference between the correlation when we use all the 64 channels and when we use a reduced number of channels. We can see that this effect is indeed consistently present for all subjects when we use a number of channels between 19 and 57. This behaviour can be seen more clearly in Fig. 4a, where the percentage of subjects with a correlation greater or equal to 100%, 95% and 90% of the correlation obtained using all the channels (green, purple and cyan lines, respectively) is shown. Fig. 4a clearly shows that for 98% of the subjects it is possible to reduce the number of channels to 19 and still obtain a correlation higher than the one obtained using all the channels. Even if we go all the way down to 8 channels, we can see that 82%, 91% and 96% of the subjects are still able to get a correlation higher than 100%, 95% and 90% of the correlation obtained using all channels, respectively.

**Fig 4.**
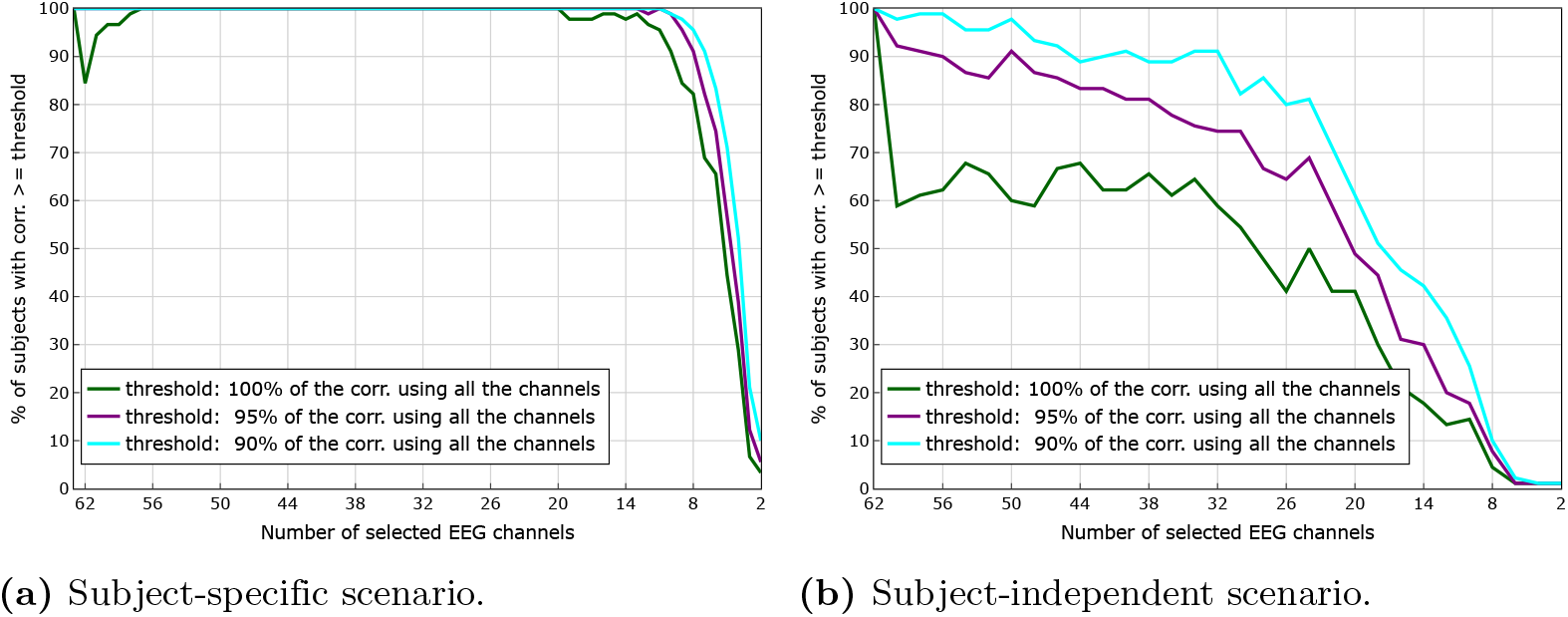
Percentage of subjects with a correlation greater or equal to 100%, 95% and 90% of the correlation obtained using all the channels. In the subject-specific scenario we can see that for 98% of the subjects is possible to reduce the number of channels to 19 and still be able to obtain a correlation higher than the one obtained using all the channels. In the subject-independent scenario we can see that for 59%, 74% and 91% of the subjects is possible to reduce the number of channels to 32 and still be able to obtain a correlation higher than 100%, 95% and 90% of the correlation obtained using all channels, respectively. The percentage of subjects can increase to 62%, 81% and 91%, respectively, if we increase the number of channels from 32 to 40.

Fig. 3d shows a comparison of the correlation obtained using the optimal number of channels (obtained through the utility metric) versus the correlation obtained using all 64 channels. In this figure we can see that for every subject we get higher correlations for the reduced number of channels, as selected by the utility metric, compared to using all channels. A Wilcoxon signed rank test showed that there was a significant difference (W=0, *p* < 0.001) between the correlation using the optimal number of channels according to the utility metric (median=0.22) compared to the one obtained using all the channels (median=0.17), which is a 29% improvement.

While for each of the 90 subjects electrodes are selected individually, we conducted some extra analysis to investigate to what extent the selected electrodes correspond across subjects. The median rank order of electrodes selected across subjects is shown in S1 Appendix Fig. 6a. While some patterns emerge, there is clearly substantial variation across subjects. In addition we analysed the correlations obtained for each subject and electrode with the forward model (see S1 Appendix Fig. 7). We found generally similar areas of interest, but there are also substantial differences. This is no surprise: in the backward model a spatial filter is designed that exploits dependencies across the channels, which is not possible in the forward model. Furthermore, in the forward model channels with highly similar information are both highlighted while one of them would be removed by the utility metric.

In the appendix the same analysis is conducted for the theta frequency band. Generally the same trends are observed as in the delta band. A Wilcoxon signed rank test showed that there was a significant difference (W=0, *p* < 0.001) between the correlation using the optimal number of channels according to the utility metric (median=0.12) compared to the one obtained using all the channels (median=0.06), which is a 100% improvement. This suggests that these results are robust to the choice of frequency band and filter parameters. However, other electrodes are selected.

#### Subject-independent electrode locations

We now consider the case where the same set of electrodes is used for all subjects. Fig. 5a shows the correlation across subjects, computed as the median across folds followed by the median across subjects. In this figure, we can see that at least 50% (median) of the subjects exhibit a stable correlation for 22 up to 64 channels.

**Fig 5.**
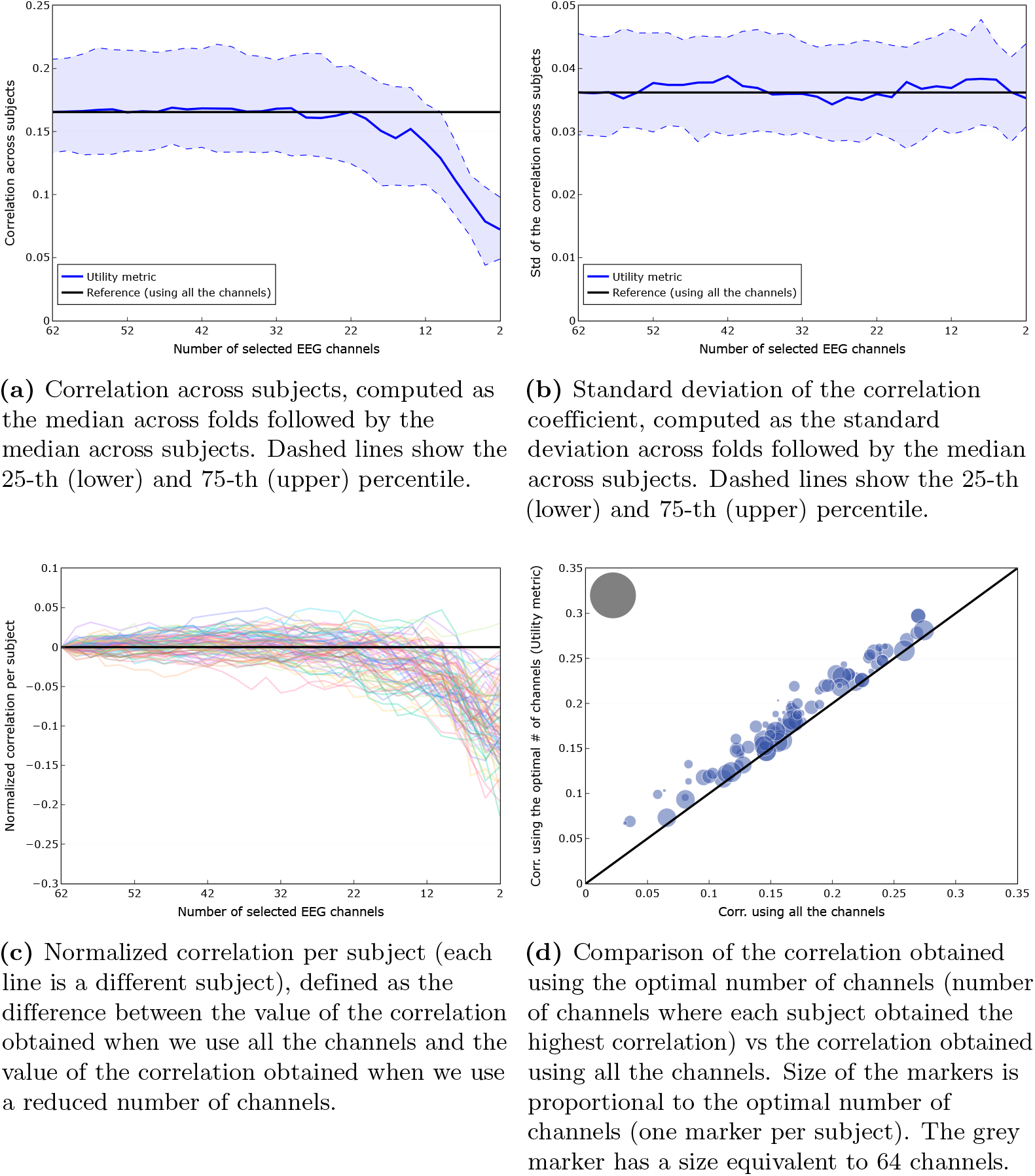
Comparison of the channel selection based on the utility metric vs using all the channels (*subject-independent scenario*). A Wilcoxon signed rank test showed that there was a significant difference (W=0, *p* < 0.001) between the correlation obtained using the optimal number of channels suggested by the utility metric (median=0.18) compared to the one obtained using all the channels (median=0.16).

Contrary to the subject-specific electrode locations, we here found a small benefit of using the symmetric channel grouping strategy: median correlations with the optimal number of channels significantly improved when moving from the channel-by-channel to symmetric grouping strategy (*W* = 1556*, p* = 0.04). In the figures and what follows, we only consider the results obtained with the symmetric grouping strategy.

Fig. 5b shows the standard deviation of the correlation, as a measure of within-subject variability, computed as the standard deviation across folds followed by the median across subjects. In this figure we can see a largely stable standard deviation of the correlation around the reference value (standard deviation of the correlation when using all the 64 channels).

Figs. 5a and 5b suggest that we could obtain a correlation value similar to the one obtained using all the available channels even if we use a reduced number of channels. However, these are group results. Fig. 5c shows, separately for each subject, the difference between the value of the correlation when we use all the 64 channels and the value of the correlation when we use a reduced number of channels. We can see that this effect is not consistently present for all subjects (if that would have been the case, all the lines would have appeared above 0 when we use a reduced number of channels *n*_*k*_, 22 ≤ *n*_*k*_ < 64). Nevertheless, a certain percentage of subjects do exhibit a higher value of the correlation when using a reduced number of channels. Fig. 4b helps us to quantify this property, by showing the percentage of subjects with a correlation greater or equal to 100%, 95% and 90% of the correlation obtained using all the channels (green, purple and cyan lines, respectively). In this figure we can see that for 59%, 74% and 91% of the subjects it is possible to reduce the number of channels to 32 and still be able to obtain a correlation higher than 100%, 95% and 90% of the correlation obtained using all channels, respectively. The percentage of subjects can increase to 62%, 81% and 91%, respectively, if we increase the number of channels from 32 to 40.

Fig. 5d shows a comparison of the correlation obtained using the optimal number of channels suggested by the utility metric versus the correlation obtained using all 64 channels. In this figure we can see that, similar to the subject-specific scenario, the utility metric consistently obtained, for every subject, a higher correlation compared to using all the channels. A Wilcoxon signed rank test showed that there was a significant difference (W=0, *p* < 0.001) between the correlation obtained using the optimal number of channels suggested by the utility metric (median=0.18) compared to the one obtained using all the channels (median=0.16).

Figs. 6a, 6b, 6c and 6d show the best 8, 16, 24 and 32 channels selected by the utility metric. Next to each group of channels (formed exactly by two electrodes, see Fig. 1), its corresponding rank order is shown, which is computed as *N* − *p* + 1, where *N* is the total number of groups and *p* is the iteration at which the group was discarded in the greedy removal procedure. The rank order reflects the importance of a group of channels with respect to the other selected groups. The lower this number, the more important the group, as it was retained for a larger number of iterations in the backwards greedy removal process due to its high influence in the LS cost (see Section). As we can see, the selected channels are mostly clustered over the left and right temporal lobes, which agrees with the empirical evidence which suggests that channels located close to auditory cortex are important for picking up electrical brain activity evoked as response to an auditory stimulus.

**Fig 6.**
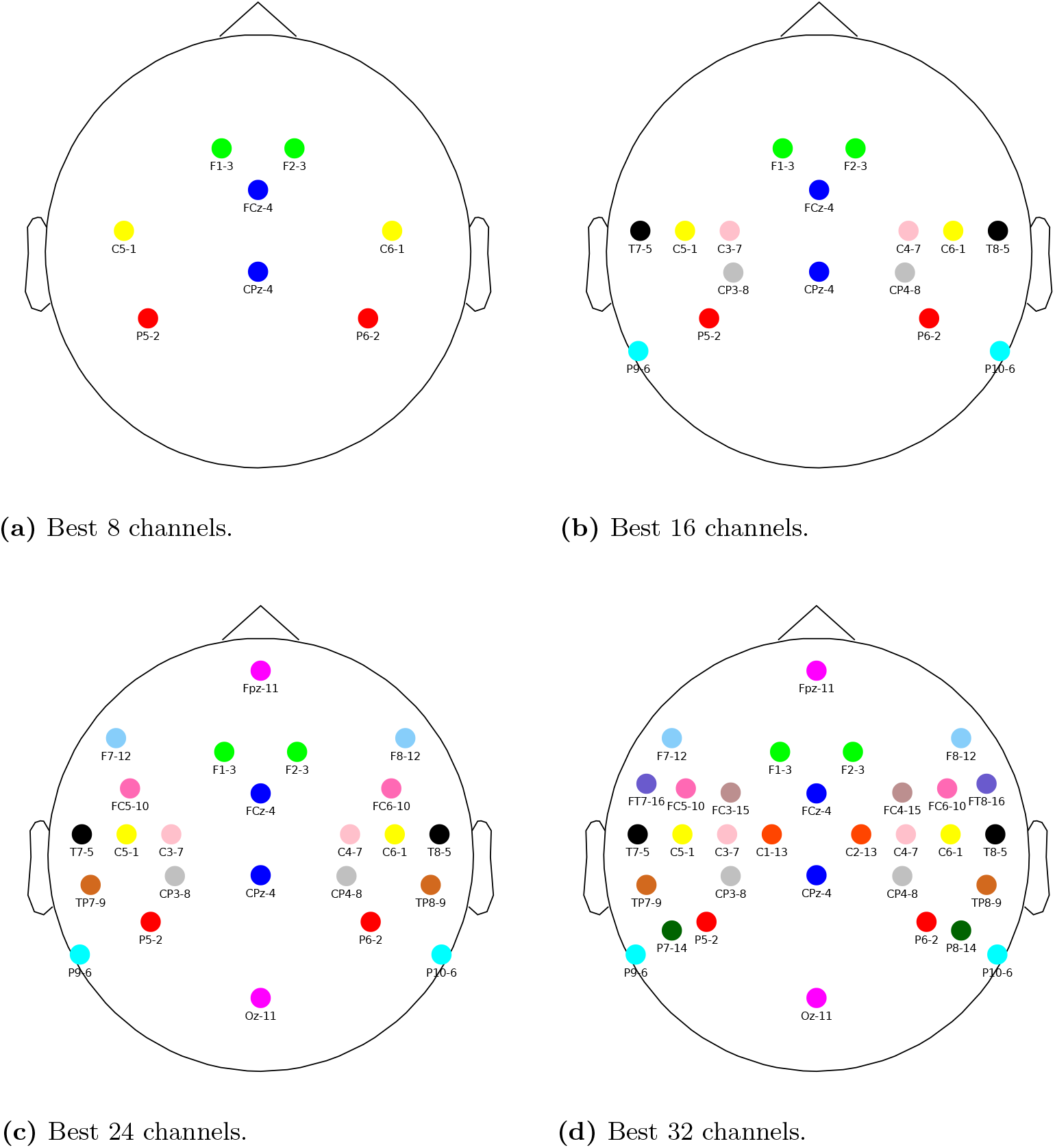
Optimal channel selection. The number next to each group of channels (formed by two electrodes, see Fig. 1) indicates the ranking of the group with respect to its influence on the LS cost (see text). The lower this number, the more important the group.

In S1 Appendix Fig. 5 the same analysis is shown for the theta frequency band. It can be seen that now mainly electrodes in the frontal and temporal areas are selected, indicating that an application (frequency-band) specific electrode selection is required.

While the electrode layouts shown in figures 6 and S1 Appendix Fig. 5 are the best we can do, they yield substantially worse performance than individual channel selections. In the 8-channel cases the percentage of subjects with unchanged performance is down to 10%, so this layout cannot be recommended for practical applications. In S1 Appendix Fig. 8 and 9 we show for each channel in each subject-independent layout how many times the same electrode was selected in the corresponding subject-specific layouts. The ranges are 20-28; 31-40; 37-55; 44-67 out of 90 subjects for respectively the 8, 16, 24 and 32 channel layout. These relatively small proportions indicate again that the generic layouts are not optimal. Note that in this analysis we did not apply the symmetric grouping constraint in the subject-independent case for the sake of comparison.

## Discussion

Based on 64-channel EEG recordings, we determined the effect of reducing the number of available channels and the optimal electrode locations on the scalp for 4 frequently-used numbers of channels. This was based on a utility-based metric, by which we avoided the computationally intractable number of combinations that underlies the problem at hand.

Mirkovic et al. (2015); Fuglsang et al. (2017) tackled the channel subset selection problem in the context auditory attention decoding (identify the attended speech stream in a multi-speaker scenario). They processed EEG recordings from 12 and 29 subjects, acquired using an EEG system with 96 and 64 channels, respectively. They found that, on average, the decoding accuracy dropped when using a number of channels less than 25. Both studies used the same channel selection strategy, which is based on an iterative backward elimination approach, where at each iteration, the channel with the lowest average decoder coefficient is removed from the next iteration. This strategy assumes that important channels will have a large coefficient in the LS solution. However, as explained in the introduction, this is not necessarily a suitable assumption. They did not report optimal electrode positions.

Narayanan and Bertrand (2019) also analyzed the channel subset selection problem in the context of auditory attention decoding, using a channel selection strategy based on the same utility metric discussed in the present study, but without imposing the symmetric grouping approach discussed in Section. They found that, on average, the decoding accuracy remained stable when using a number of channels greater or equal to 10. The (asymmetric) channels reported in their study correspond with the ones reported in this study in the sense that mostly channels around the left and right temporal lobes were selected.

Instead of attention decoding accuracy, we assessed the correlation between actual and reconstructed envelope (in a single-speaker scenario), which can be used as a metric for speech intelligibility (Vanthornhout et al., 2018; Lesenfants et al., 2019). A major difference with auditory attention decoding accuracy as a metric is that in the attention decoding paradigm correlations are compared between the attended and unattended speaker, therefore if both increase or decrease in the same direction, decoding accuracy is not affected. For subject-specific electrode locations, we found similar differences between the DMB and utility metric: using the DMB metric, on average 14 electrodes were required to avoid a drop in correlation below the 64-channel case, and using the utility metric, only 6 electrodes were required. On top of this, we found a substantial increase in correlation when reducing the number of electrodes from 64 to 32-20. This indicates that application of the proposed channel selection approach may be practically useful.

The stable or sometimes even improved performance after reducing the number of channels could be attributed to the removal of noisy or irrelevant channels that do not contribute significantly to the reconstruction of the target speech envelope. As explained in Section, the backward problem is usually solved by using a regularized Ridge regression approach, which shrinks the magnitude of many decoder components to prevent overfitting (finding solutions that minimize the reconstruction error while satisfying, at the same time, the condition of having a small norm value). We recalculated the optimal regularization parameter for each number of channels. Reducing the number of channels has a similar regularization effect; it reduces the degrees of freedom by discarding irrelevant channels, making the model less prone to overfitting.

In the case where the same channels were selected for all subjects, the initial increase in correlation with decreasing number of channels was smaller and not present for all subjects. Therefore in this case our strategy is not useful to increase correlation.

Grouping electrodes across hemispheres to remove bias effects improved the results in the subject-independent but not in the subject-specific experiments. This is probably because of subject specific hemisphere bias.

### Selected channels

Based on the literature, we expect that most of the signals of interest originate from auditory cortex (e.g., Brodbeck et al., 2018; Pasley et al., 2012). We indeed see that channels that cover dipoles originating in this area are always selected with high priority. For higher numbers of channels, other areas are covered where auditory related responses have been shown to originate from, such as the inferior frontal cortex and the premotor cortex (Das et al., 2018; Lesenfants et al., 2019), and possibly channels that aid in the suppression of large irrelevant sources.

When changing from the delta to theta frequency band, different channels were selected. This suggests differences in neural sources and indicates that the frequency band of interest should be taken into account when selecting channels.

Comparison of the selected channels with the literature is hard due to methodological differences: (Mirkovic et al., 2016) investigated the attention decoding paradigm in a two-speaker scenario (instead of single-speaker correlation in this work) and only investigated a number of a priori defined channel selections. They report high decoding accuracy using the scalp temporal and the scalp wide layouts, which is in agreement with our results.

While the presented channel layouts for 8, 16, 24, and 32 channels are the best we can do with our current data and methods, and may be useful for some applications, it should be pointed out that they yield relatively poor performance compared to subject-specific layouts and are therefore certainly not optimal for all subjects.

### Applications

The use of subject-specific or subject-independent electrode locations leads to different applications.

Subject-independent electrode locations could be used to design a headset for a specific application. For example the backward model has been proposed in applications where an objective measure of speech intelligibility is needed. Our suggested electrode positions could be used to configure an electrode cap or headset for this specific application. We chose to run our calculations with the speech envelope as the stimulus feature and for the delta band (0.5-4Hz), as these parameters are most commonly used. Note that when deviating from these parameters, the selection should be re-run. In particular, when higher-order stimulus features are used, we expect significant changes in topography and therefore optimal electrode positions.

Subject-specific electrode locations are at this point mainly useful to increase correlations when a full electrode cap is available. In this case, the utility-based algorithm would be part of the processing pipeline to retain the optimal number of electrodes. In the future subject-specific locations may also be useful to design a subject-specific headset based on initial recordings with a full cap. However, this will require validation of generalisability between test sessions and between EEG systems, which is currently unknown.

## Conclusion

In this work, the effect of selecting a reduced number of EEG channels was investigated within the context of the stimulus reconstruction task. We proposed a utility-based greedy channel selection strategy, aiming to induce the selection of symmetric EEG channel groups. We evaluated our approach using 64-channel EEG data from 90 subjects. When using individual electrode selections for each subject, we found that the correlation between the actual and reconstructed envelope first increased with decreasing number of electrodes, with an optimum at around 20 electrodes. This means that the proposed method can be used in practice to obtain higher correlations. When using a generic electrode placement (the same for all subjects), we obtained a stable decoding performance when using all 64 channels down to 22, suggesting that it is possible to get an acceptable reconstruction of the speech envelope from a reduced number of EEG channels.

## Acknowledgments

The authors would like to thank to Abhijith Mundanad Narayanan for sharing his code to compute the utility metric, as well as for the insightful discussions about the mathematical properties of the utility metric.

## Supporting information

**S1 appendix** Extra analyses

## Notes

### Competing Interest Statement

The authors have declared no competing interest.

