## Supporting information for "Effect of number and placement of EEG electrodes on measurement of neural tracking of speech"

### S1 - Appendix

#### 1 Analysis in the theta band (4-8 Hz)

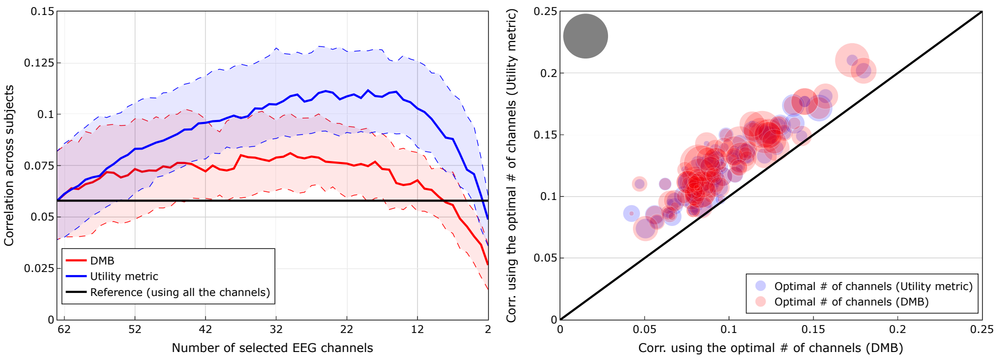


1

**(a)** Correlation across subjects, computed as the median across folds followed by the median across subjects. Dashed lines show the 25-th (lower) and 75-th (upper) percentile.

**(b)** Comparison of the correlation obtained using the optimal number of channels (number of channels where each subject obtained the highest correlation). Size of the markers is proportional to the optimal number of channels (one marker per subject). The grey marker has a size equivalent to 64 channels.

**Fig 1. Comparison of channel selection strategies: utility metric vs DMB (*subject-specific scenario* in the theta band).** A Wilcoxon signed rank test showed that there was a significant difference (W=0, $p < 0.001$) between the correlation obtained using the optimal number of channels according to the utility metric (median=0.12) compared to the one obtained using DMB (median=0.09). Another Wilcoxon signed rank test showed that there was also a significant difference (W=758.5, $p < 0.001$) between the optimal number of channels selected by the utility metric (median=20) compared to the one selected by DMB (median=29).

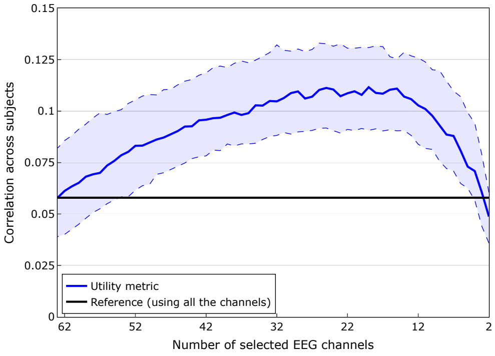


**(a)** Correlation computed as the median across folds followed by the median across subjects. Dashed lines show the 25-th (lower) and 75-th (upper) percentile.

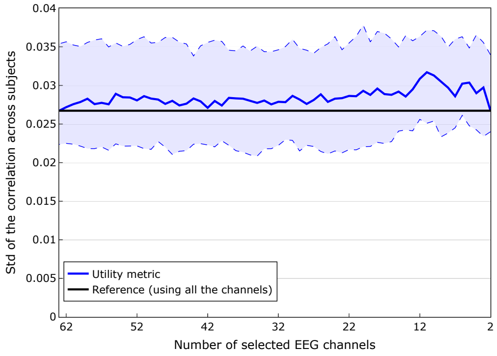


**(b)** Standard deviation of the correlation coefficient, computed as the standard deviation across folds followed by the median across subjects. Dashed lines show the 25-th (lower) and 75-th (upper) percentile.

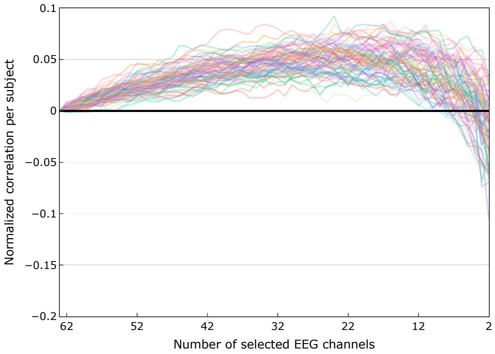


**(c)** Normalized correlation per subject (each line is a different subject), defined as the difference between the value of the correlation obtained when we use all the channels and the value of the correlation obtained when we use a reduced number of channels. For the best electrode selection, correlations were on average 98% higher than when using all the available electrodes.

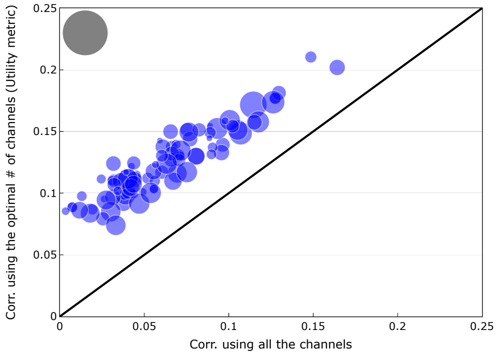


**(d)** Comparison of the correlation obtained using the optimal number of channels (number of channels where each subject obtained the highest correlation) vs the correlation obtained using all the channels. Size of the markers is proportional to the optimal number of channels (one marker per subject). The grey marker has a size equivalent to 64 channels.

**Fig 2. Comparison of the channel selection based on the utility metric vs using all the channels (*subject-specific scenario* in the theta band).** A Wilcoxon signed rank test showed that there was a significant difference (W=0, $p < 0.001$) between the correlation obtained using the optimal number of channels suggested by the utility metric (median=0.12) compared to the one obtained using all the channels (median=0.06).

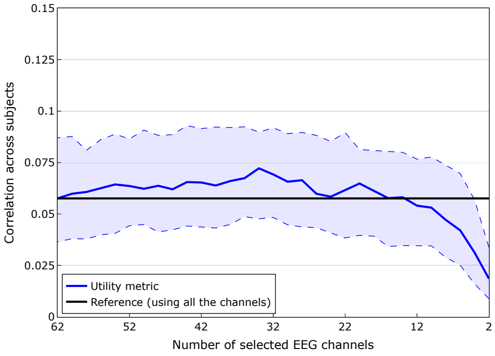


**(a)** Correlation across subjects, computed as the median across folds followed by the median across subjects. Dashed lines show the 25-th (lower) and 75-th (upper) percentile.

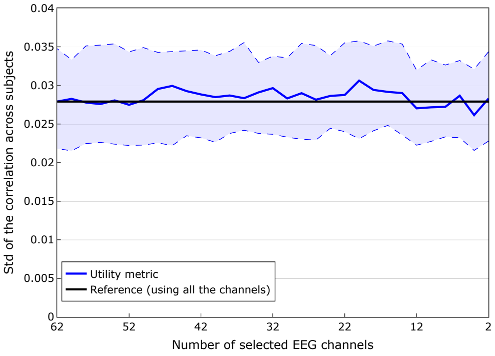


**(b)** Standard deviation of the correlation coefficient, computed as the standard deviation across folds followed by the median across subjects. Dashed lines show the 25-th (lower) and 75-th (upper) percentile.

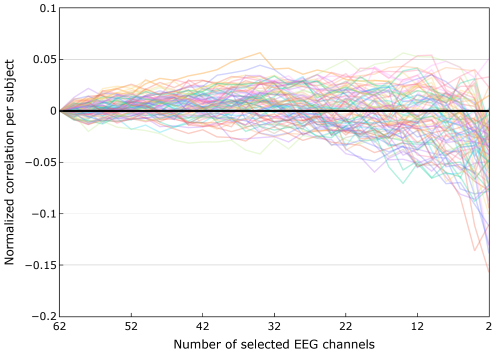


**(c)** Normalized correlation per subject (each line is a different subject), defined as the difference between the value of the correlation obtained when we use all the channels and the value of the correlation obtained when we use a reduced number of channels.

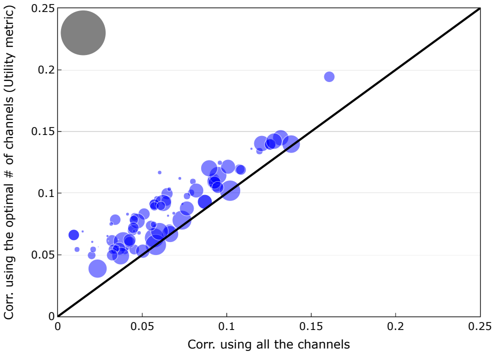


**(d)** Comparison of the correlation obtained using the optimal number of channels (number of channels where each subject obtained the highest correlation) vs the correlation obtained using all the channels. Size of the markers is proportional to the optimal number of channels (one marker per subject). The grey marker has a size equivalent to 64 channels.

**Fig 3. Comparison of the channel selection based on the utility metric vs using all the channels (*subject-independent scenario* in the theta band).** A Wilcoxon signed rank test showed that there was a significant difference (W=0, $p < 0.001$) between the correlation obtained using the optimal number of channels suggested by the utility metric (median=0.08) compared to the one obtained using all the channels (median=0.06).

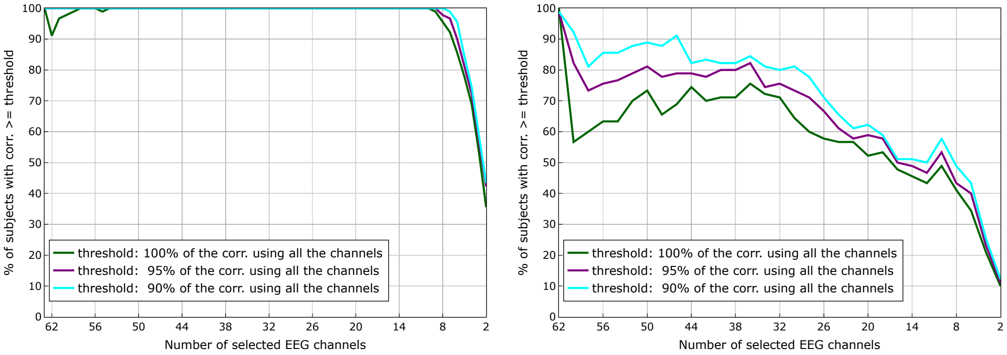


**(a)** Subject-specific scenario. **(b)** Subject-independent scenario.

**Fig 4. Percentage of subjects with a correlation greater or equal to 100%, 95% and 90% of the correlation obtained using all the channels in the theta band**. In the subject-specific scenario we can see that for 99% of the subjects is possible to reduce the number of channels to 9 and still be able to obtain a correlation higher than the one obtained using all the channels. In the subject-independent scenario we can see that for 71%, 76% and 80% of the subjects is possible to reduce the number of channels to 32 and still be able to obtain a correlation higher than 100%, 95% and 90% of the correlation obtained using all channels, respectively. The percentage of subjects can increase to 76%, 82% and 84%, respectively, if we increase the number of channels from 32 to 36.

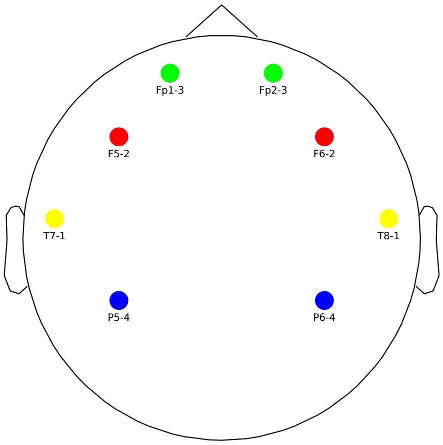


**(a)** Best 8 channels.

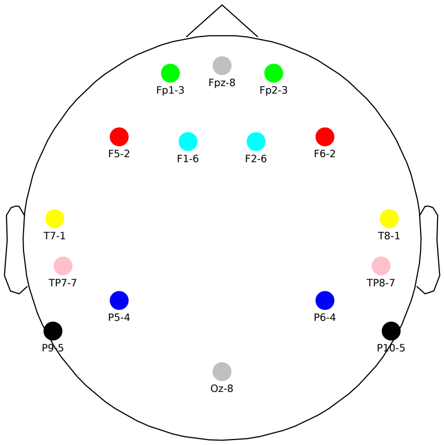


**(b)** Best 16 channels.

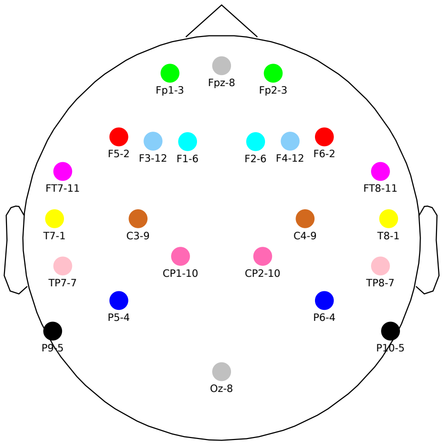


**(c)** Best 24 channels.

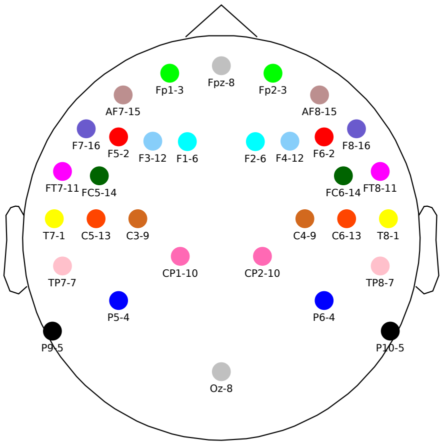


**(d)** Best 32 channels.

**Fig 5. Optimal channel selection for the theta band.** The number next to each group of channels (formed by two electrodes, see Figure **??**) indicates the ranking of the group with respect to its influence on the LS cost (see text). The lower this number, the more important the group.

#### 2 Additional subject-specific analysis

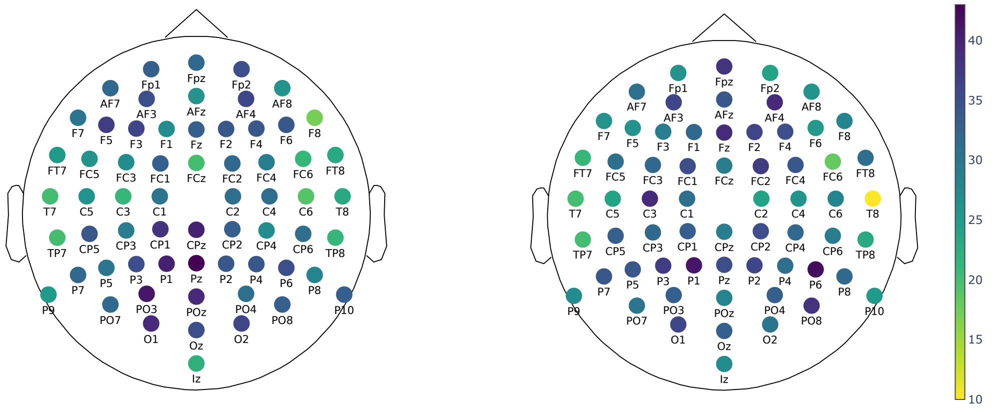


**(a)** Delta. **(b)** Theta.

**Fig 6. Median rank order across subjects of each channel, subject-specific scenario, utility metric.** The rank order of a channel, provided by the utility metric, reflects the importance of a channel with respect to the other selected channels. The lower the rank, the more important the channel.

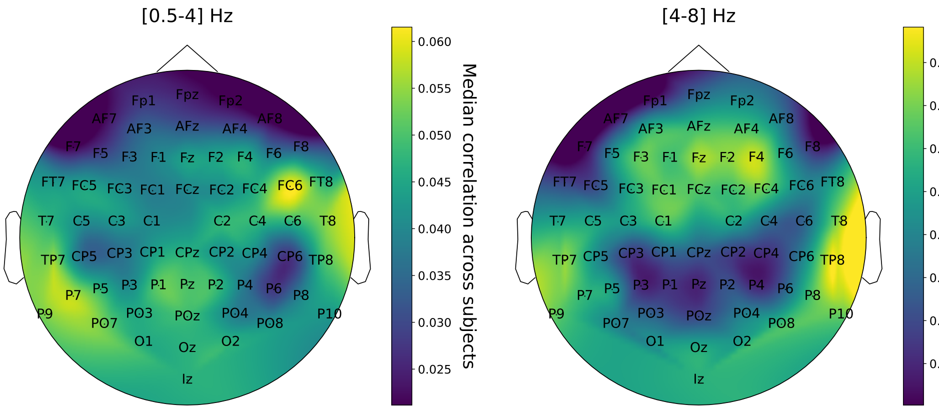


**Fig 7. Median correlation for the forward model, subject-specific scenario.** The forward model was calculated for each subject and electrode, and the median correlation between actual and predicted EEG is plotted per electrode. By comparing with Figure 6, we can see that for both the delta and theta bands, channels with lower rank order (more important channels) are generally also channels with higher correlation in the forward model. For the delta band such channels are primarily concentrated in the temporal and pre-frontal regions, whereas for the theta band they are located in the temporal and frontal regions. These locations agree with the optimal channel selection for both the delta and theta band.

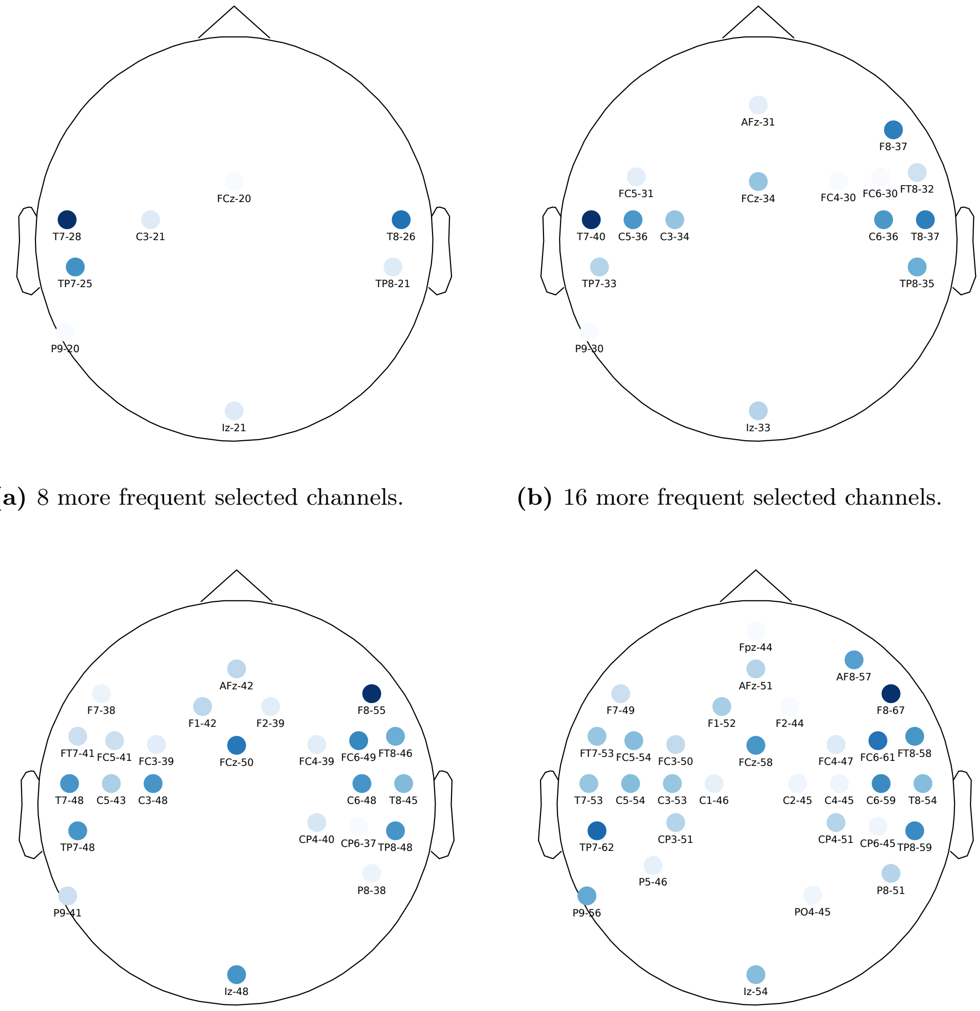


**(a)** 8 more frequent selected channels.

**(b)** 16 more frequent selected channels.

**(c)** 24 more frequent selected channels.

**(d)** 32 more frequent selected channels.

**Fig 8. Channel selection frequency (Delta band).** The color indicates for each subject-independent layout (without symmetric grouping constraint) for how many subjects each channel was selected in the subject-specific case. The number next to the channel label indicates the number of subjects for whom the channel was selected out of the total number of 90 subjects.

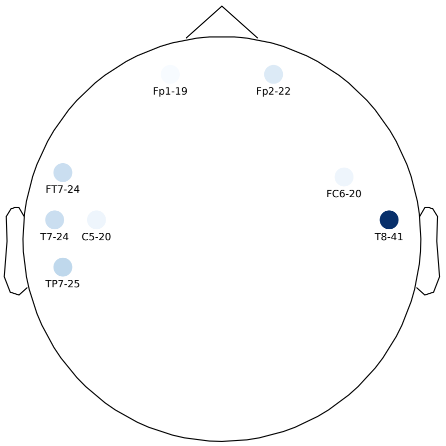


**(a)** 8 more frequent selected channels.

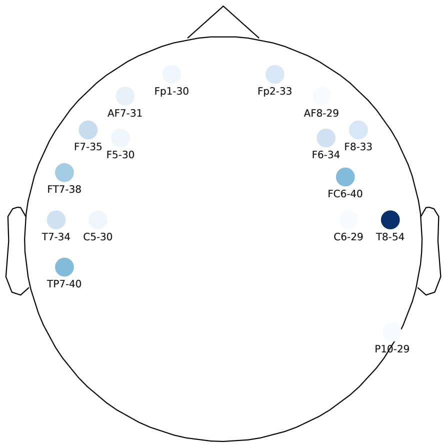


**(b)** 16 more frequent selected channels.

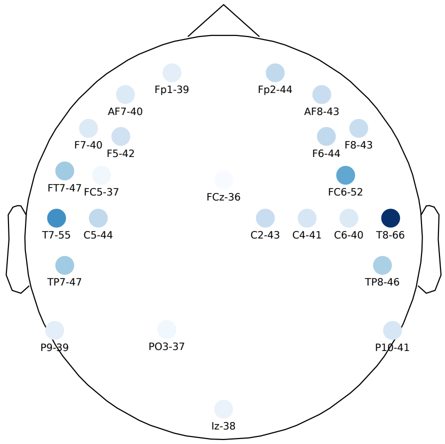


**(c)** 24 more frequent selected channels.

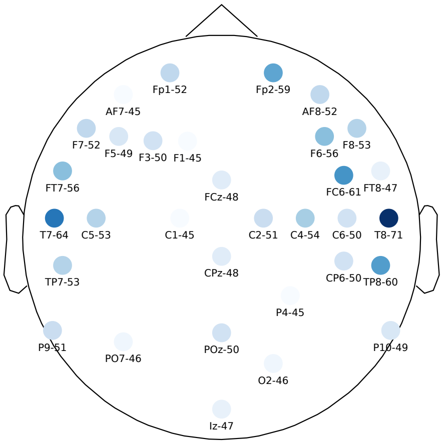


**(d)** 32 more frequent selected channels.

**Fig 9. Channel selection frequency (Theta band).** The color indicates for each subject-independent layout (without symmetric grouping constraint) for how many subjects each channel was selected in the subject-specific case. The number next to the channel label indicates the number of subjects for whom the channel was selected out of the total number of 90 subjects.
